# On the global attractors in one mathematical model of antiviral immunity

**DOI:** 10.1101/2020.10.16.343145

**Authors:** Alexei Tsygvintsev

## Abstract

We consider the mathematical model introduced by Batholdy *et al.* [1] describing the interaction between viral pathogens and immune system. We prove the global asymptotic stability of the infection steady-state if the basic reproductive ratio *R*_0_ is greater than unity. That solves the conjecture announced in [7].

## 1. Introduction

Our immune system is a complex multifunctional mechanism aimed to protect the host from various external infectious pathogens (viruses, bacteria etc). It is well known that the emergence and development of numerous diseases is largely controlled by the immune defence factors, and therefore their study and modelling is of great biological and medical interest. The most important players in our adaptive immune system are the B-lymphocytes [4]. These are white blood cells whose function is to produce antibodies - the *Y*-shaped special proteins that bind to specific antigens tagging them to be eliminated. The role of T-lymphocytes is very different [4]. They directly eliminate pathogenic microorganisms.

The human immune defence has been developed over thousands of years and we are still far from complete understanding all the complex mechanisms involved.

As we explore the mechanisms of pathogen interactions with the immune system, a number of important simplifications have to be employed. In the mathematical modelling it is often necessary to consider as a single group the different immune system participants. At the same time, the known mechanisms of cellular interactions should be simplified in the hope that the most important biological features are satisfactory reflected.

Last years, various models have been proposed in attempts to understand better the interaction between the immune system and viral pathogens [10]. Two principal types of immune response to viral intrusion can be underlined: *lytic* (destroying directly infected cells) and *nonlytic* (inhibiting the viral replication by slowing it down) [9]. In the next section we will consider one of such models describing the interaction of immune lytic/nonlytic factors with viral pathogens.

## 2. The model of antiviral immunity

In 2000 Bartholdy *et al.* [1] proposed a mathematical model consisting of 3 ordinary differential equations to describe the outcome of the competition between the virus and the human immune system. Below we describe briefly these equations.

Let *x*(*t*) be the number of uninfected host cells, *y*(*t*) is the number of cells infected by the virus at the moment *t* and *z*(*t*) is the number of immune cells population. Usually, the values *x*(*t*), *y*(*t*), *z*(*t*) are considered as concentrations, i.e. the number of cells per unit of volume.

To simplify the model it is assumed that the dynamics of the free virus circulating in the blood is not directly taken into account i.e. only already infected cells *y*(*t*) are considered. This is based on the realistic assumption that the rate of the free virus evolution is much faster than that of already infected cells [9]. The case where the free virus dynamics is taken into account has been considered in the work [6].

The first equation of the model describes the evolution of uninfected host cells:

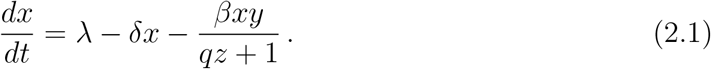

Here λ is the production rate of host cells, *δx* is the death rate and *βxy* is the infection rate once the immune system is not active. The term *qz* + 1 is the nonlytic inhibition rate.

The second equation controls the dynamics of the population of infected cells:

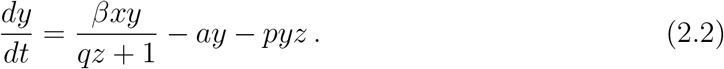

The infected cells die naturally with the rate *a* and are killed by the lytic immune response with the rate *pyz*. Finally, the third equation governs the evolution of immune cells population:

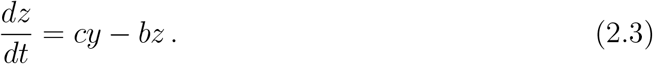

The immune cells are produced with the rate *cy* and naturally disappear with the decay rate *bz*. We assume in this paper that all parameters of the model (2.1)–(2.3) are strictly positive real numbers.

As shown in [7], every solution of the differential equations (2.1)–(2.3) with initial conditions *x*(0), *y*(0), *z*(0) ≥ 0 is bounded and verifies *x*(*t*), *y*(*t*), *z*(*t*) ≥ 0 for all *t* ≥ 0.

## 3. Steady states and their stability

The asymptotic behaviour of solutions of the model (2.1)–(2.3) is of great interest. In biology and medicine, processes often go through an intermediate stage before reaching the established state called an *attractor*. In the simplest case, the attractor can be a steady-state i.e. the equilibrium point (*x*_0_, *y*_0_, *a*_0_) at which the velocity vector vanishes: *dx*/*dt* = *dy*/*dt* = *dz*/*dt* = 0. In a general situation, it can be a periodic solution or a set with a very complex internal structure, as in the Lorenz case [8].

The steady-state can be stable or unstable. In the stable case, all neighbourhood trajectories converge to it asymptotically and in the unstable case some of them can be repelled. More precisely, the following definition of global asymptotic stability will be used in this paper (see [2]):

### Definition 3.1.

*Let U* ⊂ ℝ^*n*^ *be an open set and X*′ = *V* (*X*), *X* ∈ *U be a system of n differential equations where the corresponding vector field V is continuous and such that all its solutions t* ⟼ *X*(*t*) *extend* ∀*t* ∈ ℝ^+^. *The equilibrium X*_0_ ∈ *U is globally asymptotically stable iff* ||*X*(*t*) − *X*_0_|| → 0 *as t* → +∞ *for any solution with X*(0) ∈ *U i.e. the basin of attraction of X*_0_ *is the whole set U*.

As already noticed in works [1], [9], the asymptotic behaviour of solutions of the system (2.1)–(2.3) largely depends on the value of the basic reproductive ratio 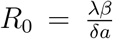. By definition, this key biological parameter describes the average number of newly infected cells generated from one infected cell at the beginning of the infectious process. This is a fundamental characteristic that determines whether a virus develops within the host or it is eliminated finally by the immune response.

Depending on the value of *R*_0_, the model (2.1)–(2.3) can exhibit two different kinds of steady-states described below.

The infection-free state is characterised by the property that *y* = 0 i.e. the virus was completely cleared by the immune system. The following result was shown in [7] using the Lyapunov’s method [5]:

### Theorem 3.1.

Let *R*_0_ ≤ 1, *then the only equilibrium of the* *equations* (2.1)–(2.3) *contained in the domain x*, *y*, *z* ≥ 0 *is infection-free and given by*

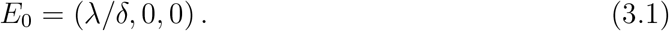

*Moreover it is globally asymptotically stable in the domain x*, *y*, *z* ≥ 0.

We provide below an independent simple argument to show that *y*(*t*) → 0 as *t* → +∞ if *R*_0_ < 1.

Using the equations (2.1)–(2.3) one finds by derivation:

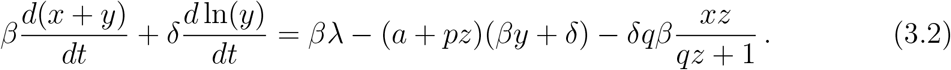

Since all quantities *x*,*y*,*z* are positive, the last equality implies that

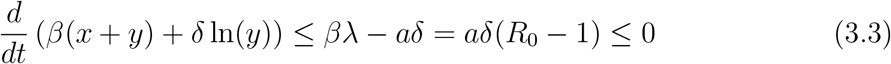

Let us assume that *R*_0_ < 1. Then, integrating the previous inequality over the interval [0, *t*], *t* > 0, we find

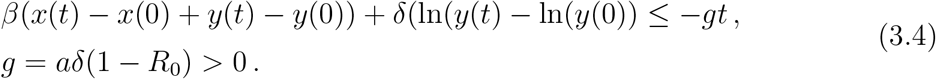

Thus ln(*y*(*t*)) → −∞ as *t* → +∞ since *x*(*t*), *y*(*t*) ≥ 0, ∀ *t* ≥ 0. But it yields immediately that

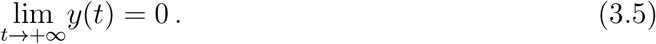

Now let us turn our attention to the situation *R*_0_ > 1. Here, the disease can finally develop within the host since the strength of immune response is not strong enough to clear the infection. The next proposition shows that in this case there exits unique infection steady-state (*x*_1_, *y*_1_, *z*_1_) with *y*_1_ > 0 and provides its simple rational parametrisation which is new and was not considered previously to the best of our knowledge.

### Proposition 3.1.

*Let R*_0_ > 1, *then the only infection steady-state of the model* (2.1)–(2.3), *contained in the domain x*, *y*, *z* ≥ 0, *is given by*

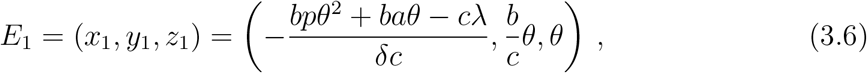

*where θ* ∈ (0, *θ*_0_), 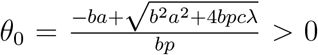 *is the parameter defined (uniquely) from the following equation*:

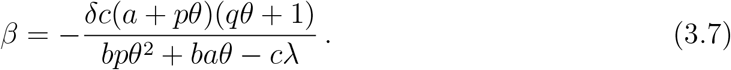

*Proof*. The proof is straightforward and uses the definition of the equilibrium point *dx*/*dt* = *dy*/*dt* = *dz*/*dt* = 0.

### Remark 3.1.

*Once R*_0_ *crosses the critical value R*_0_ = 1 *(on the left), the equilibrium point E*_0_ *interchanges its local stability with E*_1_ *which appears through the transcritical bifurcation. That can be derived from the results of the work* [7] *where the local stability of E*_0,1_ *was analysed and the corresponding eigenvalues were calculated.*

Early attempts (see [7]) to prove the global asymptotic stability of *E*_1_ succeeded only for some special parameters of the system (2.1)–(2.3), namely for *q* = 0 or if *R*_0_ is sufficiently close to unity. Nevertheless, it was conjectured by the same authors that the global asymptotic stability of *E*_1_ holds for arbitrary parameter values provided that *R*_0_ > 1.

Our result below solves this conjecture.

### Theorem 3.2.

*The infection steady state E*_1_ *is globally asymptotically stable if R*_0_ > 1.

*Proof.* The proof is based on the Lyapunov’s method [5]. We will try to give a clear idea of how the suitable Lyapunov function can be found and we hope this construction will be useful in other similar problems. Let *D* be the Lie derivative operator associated to the differential equations (2.1)–(2.3). We are looking for 3 differentiable functions *I*: *U* → ℝ, *U* = ℝ^3+^* those derivatives *D*(*I*) are constant on leaves of some codimension-1 foliation of *U*. Such functions can be easily found from the system (2.1)–(2.3):

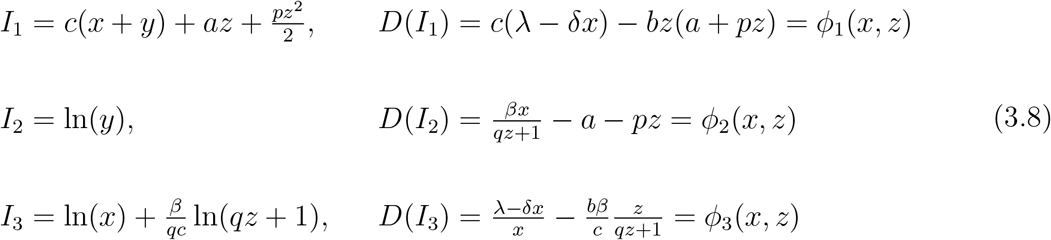

As seen from these formulas, *D*(*I_k_*), *k* = 1, 2, 3 do not depend on the variable *y* i.e. satisfy our requirement. At the same time, *ϕ_k_*(*x*_1_, *z*_1_) = 0, *k* = 1, 2, 3 since *E*_1_ = (*x*_1_, *y*_1_, *z*_1_) is the equilibrium point. We determine now the real constants *a*_1_, *a*_2_, *a*_3_ in such a way that *E*_1_ is a critical point of the Lie derivative *L*\* = *D*(*L*) where *L* is defined by

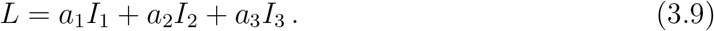

As shown later, *L* will be our Lyapunov function.

The equation grad(*L*\*)(*x*_1_, *z*_1_) = 0, up to a certain multiplier, has the unique solution for *a*_1_, *a*_2_, *a*_3_ given by *a*_1_ = −1*/c*, *a*_2_ = *y*_1_, *a*_3_ = *x*_1_. It is straightforward to check that *L* possesses *E*_1_ as the unique extremum in *U* which is the global maximum.

Let *U*_1_ = ℝ^2+^*. The next lemma guarantees the positiveness of *L*\* in *U*_1_.

### Lemma 3.1.

*The point* (*x*_1_, *z*_1_) *is the global minimum of L*\* *in U*_1_ *such that L*\*(*x*_1_, *y*_1_) = 0 *and L*\*(*x, z*) > 0 *for all* (*x, z*) ∈ *U*_1_ \ {(*x*_1_, *z*_1_}.

Thus, *L* is a Lyapunov function of (2.1)–(2.3) with *D*(*L*) ≥ 0 in *U* and *D*(*L*) = 0 along the line *I* = {(*x, y, z*) ∈ *U*: *x* = *x*_1_, *z* = *z*_1_, *y* ∈ ℝ^+^*}. By the unicity, *E*_1_ is the only invariant set contained in *I*. All solutions of (2.1)–(2.3) are bounded for *t* ≥ 0 as shown in [7]. So, according to the LaSalle’s theorem (see [3], p. 524, Theorem 3, [5]) the equilibrium point *E*_1_ is globally asymptotically stable in *U*.

Below we sketch the main lines of the proof of Lemma 3.1.

All algebraic computations are greatly simplified by the fact that *L*\* is a function of *x*, *z* only and its sign can be easily controlled. That was the main underlaying idea of how the function *L* was designed. The elementary calculation show that (*x*_1_, *z*_1_) is a local minimum of *L*\*.

The function *L*\* can be written as the ratio of two rational functions *f*, *f*_1_ depending each on *x* and *z* only:

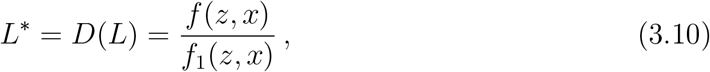

where *f*_1_(*z, x*) = *c*(*qz* + 1) > 0 in *U*_1_ and the numerator *f* can be written as

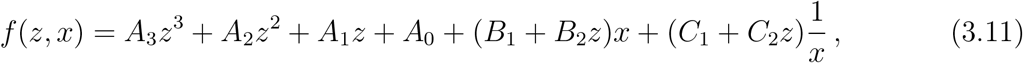

with known constants *A_i_, B_j_, C_k_* given in Appendix and depending on the parameters of the system (2.1)–(2.3). In particular, *A*_3_, *B_j_, C_k_ >* 0.

Fixing *z* ∈ ℝ^+^ and varying *x* in ℝ^+^*, one can compute the global minimum of *f* as a function of *z* only. It is given by

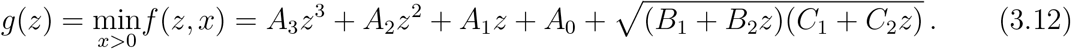

One verifies that *g*(0) > 0 using the known expressions for *A*_0_, *B*_1_ and *C*_1_. Now, finding the minimum of *L*\* in *U*_1_ can be replaced by the similar problem for *g* in ℝ^+^.

Since (*x*_1_, *z*_1_) is a critical point of *L*\* in *U*_1_ and *L*\*(*x*_1_, *z*_1_) = 0 we have *g*(*z*_1_) = *g*′(*z*_1_) = 0 where *z*_1_ = *θ* > 0 according to (3.6). One checks directly that *g*′′(*z*_1_) > 0 and so *z* = *z*_1_ is a local minimum of *g*. Using *A*_3_ > 0, by derivation with help of (3.12), one verifies that *g*′′′(*z*) > 0 for all *z* ∈ ℝ^+^ i.e. *g*′ is strictly convex and therefore it has at most 2 zeros for *z* ≥ 0. This argument, together with *g*(0) > 0 and 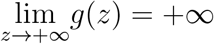, shows that *g* attains the global minimum at *z* = *z*_1_, *L* (*z*_1_) = 0 and *L* (*z*) > 0 for *z* ∈ ℝ^+^ \ {*z*_1_}. That finishes vthe proof of Lemma 3.1.

## 4. Conclusion and numerical simulations

Mathematical modelling of infectious diseases is an important tool in the development of new treatments and is necessary for a better understanding of all underlaying mechanisms of the immune defence. Our work contributes to understanding the asymptotic behaviour of solutions in one particular model (2.1)–(2.3) introduced in [1] describing the competition between lytic/nonlytic immune system and the viral pathogens. Completing the results already obtained in [7], we can finally state the existence of the sharp threshold depending on the value of the basic reproductive ratio *R*_0_. If *R*_0_ ≤ 1 then the immune system finally clears the infection i.e the steady-state where no infected cells are present is attained. If *R*_0_ > 1, as stated by Theorem 3.2, the infection develops and converges finally to the steady-state where it stays permanently controlled by the immune system. In particular, our result excludes existence of any periodic behaviour.

The system of differential equations (2.1)–(2.3) was solved numerically for the parameter values *λ* = 1, *δ* = 0.8, *q* = 10, *a* = 0.1, *p* = 5, *c* = 0.01, *b* = 1, *β* = 0.15 with the corresponding basic reproductive ratio *R*_0_ = 1.94 and *θ* = 0.01. The infection steady-state in this case, as given by formulas (3.6), is *E*_1_ = (1.06, 1, 0.01). On Figure 1 the solution *x*(*t*), *y*(*t*), *z*(*t*) with initial conditions *x*(0) = 10, *y*(0) = 30, *z*(0) = 30 is represented for the period of 10 days. The Figure 2 contains the different sections of the surface levels of the Lyapunov function *L* by the plane *z* = 0.01.

**Figure 1:**
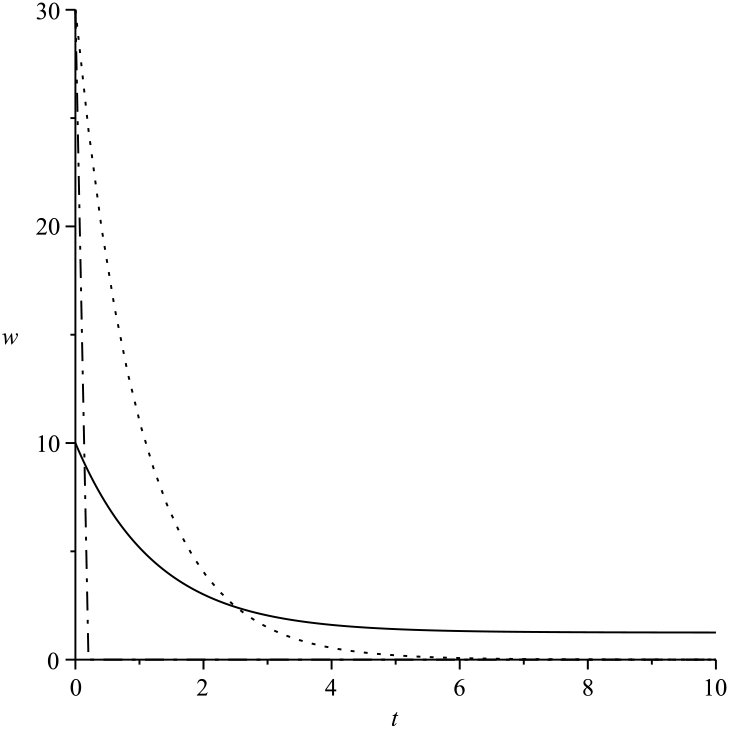
Concentration *w* of uninfected host cells *x* (solid), infected cells *y*(*t*) (dash) and the immune cells *z*(*t*) (dot) for the period of 10 days.

**Figure 2:**
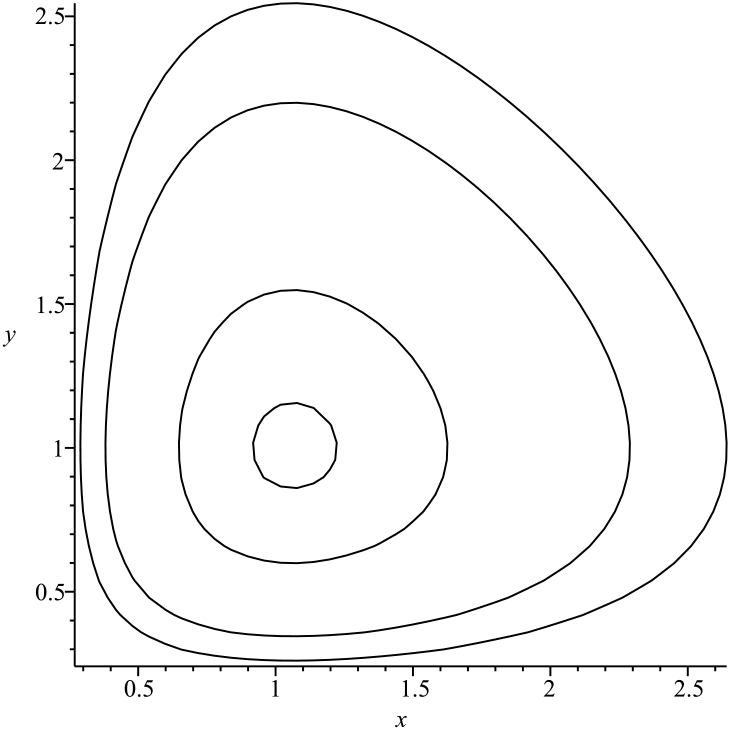
Sections by the plane *z* = 0.01 of level surfaces *L* = const for −*L* = 4.96, 4.76, 4.46, 4.36, 4.26. The corresponding value *L*(*x*_1_, *y*_1_, *z*_1_) is −1.96.

In conclusion, we will mention a number of open questions concerning the model (2.1)–(2.3). First of all, for each specific type of infectious pathogens it is necessary to develop an effective method of evaluating its parameters using experimental data.

Secondly, it is necessary to obtain accurate estimates of the time of convergence to steady states *E*_1,2_, as well as to study in more detail the qualitative behaviour of solutions during the stabilization period.

## Appendix

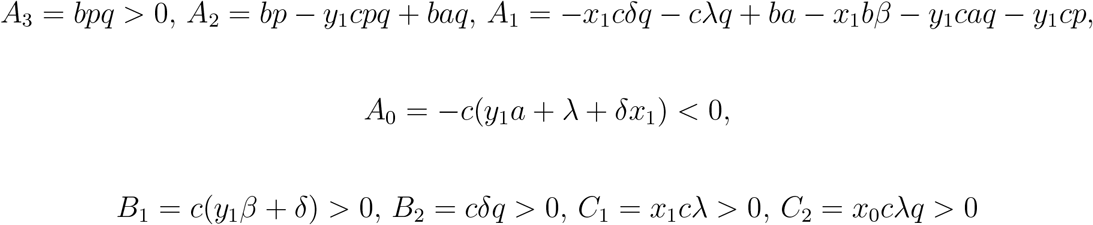

## References

[1] C. Bartholdy, J P Christensen, D. Wodarz, and A. R. Thomsen. Persistent virus infection despite chronic cytotoxic t-lymphocyte activation in gamma interferon-deficient mice infected with lymphocytic choriomeningitis virus. Journal of Virology, 74:10304–10311, 2000.

[2] P. Hartman and C. Olech. On global asymptotic stability of solutions of differential equations. Transactions of the American Mathematical Society, 104:154–178, 1962.

[3] J. LaSalle. Some extensions of liapunov’s second method. IRE Transactions on Circuit Theory, 7:520–527, 1960.

[4] K. Murphy, P. Travers, M. Walport, and C. Janeway. Janeway’s immunobiology. Garland Science, New York, 2012.

[5] J. La Salle and S. Lefschetz. Stability by Liapunov’s Direct Method: With Applications. Academic Press, New York, 1961.

[6] K. Wang, Y. Jin, and A. Fan. The effect of immune responses in viral infections: A mathematical model view. Discrete and Continuous Dynamical Systems - Series B, 19:3379–3396, 2014.

[7] K. Wang, W. Wang, and X. Liu. Global stability in a viral infection model with lytic and non-lytic immune responses. Comput. Math. Appl., 51:1593–1610, 2006.

[8] R. F. Williams. The structure of lorenz attractors. Inst. Hautes études Sci. Publ. Math., 50:73–99, 1979.

[9] D. Wodarz, J. P. Christensen, and A. R. Thomsen. The importance of lytic and non-lytic immune responses in viral infections. Trends Immunol., 23:194–200, 2002.

[10] C. Zitzmann and L. Kaderali. Mathematical analysis of viral replication dynamics and antiviral treatment strategies: From basic models to age-based multi-scale modeling. Front Microbiol., 9:1546, 2018.

